# Reparative and regenerative immature neutrophil-like population derived from HL-60 cells

**DOI:** 10.64898/2026.05.11.724223

**Authors:** Subhpreet Kaur, Ashu Shukla, Anjali Gupta, Byanjana Bashyal, Vyshak Suresh, Uma Nahar Saikia, Parul Chawla Gupta, Manni Luthra-Guptasarma

## Abstract

Unlike the conventional mature neutrophils, immature neutrophils have been investigated for their regenerative properties; however, their limited availability necessitates alternative generation strategies. Here, we used a combination of dimethylsulfoxide (DMSO) and 1α,25-dihydroxyvitamin D3 (D3) to differentiate myeloid leukemia (HL-60) cells into immature neutrophil-like cells. Differentiated cells exhibited reduced cell size, loss of uniformity, decreased nuclear-to-cytoplasmic ratio, band-shaped nuclei, increased proportion of CD11b^+^CD14^+^ cells (indicative of immature neutrophils), decreased proportion of CD11b^+^CD16^+^ cells (indicative of mature neutrophils), higher levels of arginase 1, TGFβ1 (markers of immature neutrophils), and no expression of CD16, MRC1 (markers of mature neutrophils and M2 macrophages, respectively). Proteomic analysis revealed enrichment of proteins associated with immature neutrophils and wound healing. Functionally, these cells supported limbal stem cell growth and wound closure *in vitro*, indicating relevance for corneal regeneration. Administration of these cells to *ex-vivo* and *in-vivo* alkali-injured corneas, resulted in significant effect on promotion of wound healing, with epithelial regeneration and decreased fibrotic markers, proving that such cells hold promise for clinical translation as a therapeutic tool for tissue repair.

## Introduction

Neutrophils are the most common polymorphonuclear innate leukocyte population, serving as the body’s immediate first responders to any infection or inflammation. Conventionally, they were considered to be a homogenous population of short-lived, terminally differentiated cells, with conserved physiological functions. However, recent studies (Deniset and Kubes, 2018; Liew and Kubes, 2019; Xie et al, 2020; He et al, 2025; Gysemans et al, 2025) have revealed their heterogeneity and diverse functional roles in both steady-state and pathological conditions. Based on different parameters, various neutrophil subsets have been described, including variations in maturity, buoyancy, localization, cell surface marker expression and function. In addition to their role in pathogen clearance, neutrophils are also understood to be involved in anti-tumor response as well as wound healing (Fridlender et al, 2009; Liu et al, 2023; Feng et al, 2025; Renò et al, 2025).

Immature neutrophils belong to this diverse group of neutrophils, which have gained recent attention due to their regenerative properties (Pillay et al, 2012; Deniset et al, 2017; He et al, 2025; Gysemans et al, 2025), useful in the contexts of tissue regeneration and homeostasis (Deniset et al, 2017; Sas et al, 2020; Jerome et al, 2022; Dong et al, 2024; Jerome et al, 2024; Yang et al, 2025). Immature neutrophils are being investigated in various models of infection and central nervous system injury. Unlike conventional neutrophils, these cells are alternatively activated, exhibit band shaped nuclei, and express markers such as arginase 1(Arg 1), interleukin-4receptor α-chain (IL4ra), transforming growth factor beta-1 (TGF β1), and insulin-like growth factor (IGF-1), along with CD11b^+^Ly6G^low^CD14^high^CD101^low^ phenotype.

A major limitation in studying any such neutrophil population is their low abundance in peripheral blood mononuclear cells (PBMCs) or bone marrow, coupled with difficulties in culturing them for long-term. To overcome this challenge, the promyelocytic leukemia cell line, HL-60, has been used as an alternative model, since it can be differentiated into different immune lineages using varied chemical inducers. HL-60 cells can be driven towards granulocytes by polar planar compounds such as dimethylsulfoxide (DMSO), dimethylformamide (DMF), or retinoic acid (RA), and towards monocytes/macrophages by 1α,25-dihydroxyvitamin D3 (the active form of vitamin D3, hereafter termed as D3) or phorbol esters (Fontana et al, 1981; Fibach et al, 1982; Collins et al, 1987; Birnie, 1988). Although DMSO is the most commonly used reagent for inducing granulocytic (primarily mature neutrophilic) differentiation (Rincón, 2018), attempts have also been made to differentiate HL-60 cells with DMSO to generate immature neutrophils (Newburger et al, 1979; Sas et al, 2020) by altering the duration of exposure to DMSO; however the yields are poor.

Interestingly, while retinoic acid promotes differentiation of HL-60 into granulocytes (Honma et al, 1980; Breitman et al, 1980) and 1α,25-dihydroxyvitamin D3 favours differentiation into monocytes/macrophages (Mangelsdorf et al, 1984), Takahashi et al. (2014) reported that the combination of retinoic acid and 1α,25-dihydroxyvitamin D3 resulted in a population of polarized, M2 macrophages, expressing higher levels of CD14, CD163, Arg 1 and TGFβ1 compared to either of the two inducers alone. We wished to examine alternative combinations of reagents to generate immature neutrophil-like cell population. Based on this, we investigated the combinatorial effect of DMSO and 1α,25-dihydroxyvitamin D3 to examine the possibility of generation of increased population of immune cells resembling immature neutrophils, and examined their regenerative effects on corneal wound healing using *in vitro*, *ex vivo* and an *in vivo* model.

## Results

### HL-60 cells induced with DMSO+D3 display reduced nuclear-to-cytoplasmic ratio, with cells exhibiting band-shaped nuclei

Unlike the uninduced HL-60 cell population, distinct morphological alterations were observed in HL-60 cells induced with various reagents. Uninduced cells and those induced with D3 alone, exhibited a uniform shape and size, maintaining a predominantly round morphology. In contrast, cells induced with DMSO alone or with DMSO+D3, showed loss of uniformity in cell shape and reduction in cell size, along with the formation of membrane projections, as indicated by arrows (Fig 1A). Leishman staining further revealed a larger nuclear-to-cytoplasmic ratio in uninduced HL-60 cells. In contrast, HL-60 cells induced with DMSO or D3 alone, exhibited a slightly less basophilic cytoplasm, with a decreased nuclear-to-cytoplasmic ratio. When HL-60 cells were induced with DMSO+D3, it was observed that in addition to displaying a reduced nuclear-to-cytoplasmic ratio, several of these cells exhibited band-shaped nuclei (Fig 1B).

**Figure 1.**
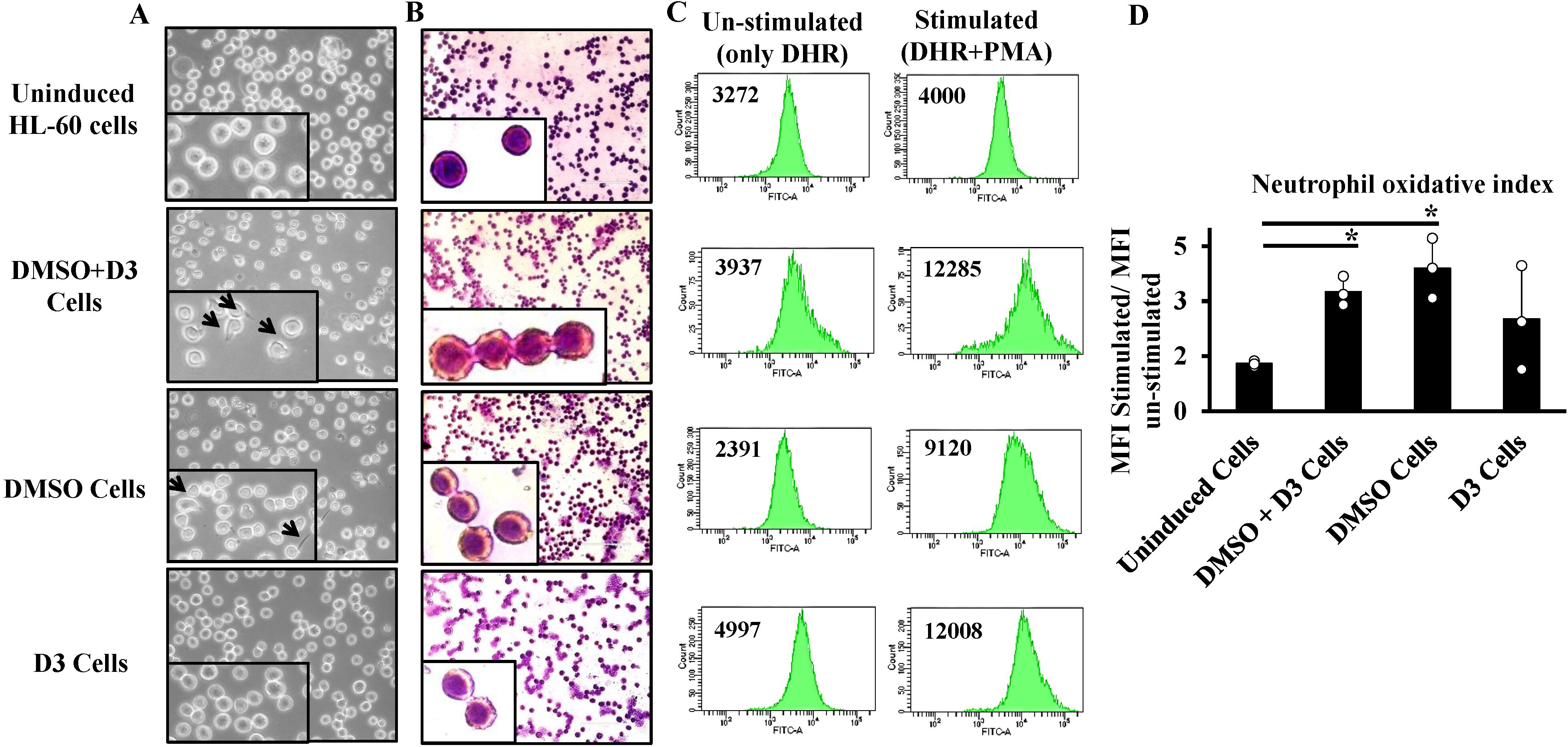
Morphological changes and reactive oxygen species (ROS) production by HL-60 cells, upon induction with different reagents. (A) Representative phase-contrast images showing morphological changes and (B) Leishman-stained images highlighting nuclear and cytoplasmic features, captured at 40X magnification (scale bar: 100 µm). (C) Flow cytometry plots for the Dihydrorhodamine (DHR) assay assessing ROS production. (D) Quantification of neutrophil oxidative index, calculated as the ratio of stimulated to un-stimulated median fluorescence intensity. Data are represented as mean ± standard deviation from three independent experiments (n=3). Statistical significance was calculated by Student’s t-test; where * denotes *P* ≤ 0.05. HL-60 cells were either uninduced or induced with DMSO+D3, DMSO alone or D3 alone for 4 days.

### DMSO+D3-induced HL-60 cells produce reactive oxygen species (ROS) upon stimulation

The DHR (Dihydrorhodamine 123) assay was used to assess the potential of ROS production by HL-60 cells upon exposure to various inducers. Cells were evaluated for their ability to oxidize the freely permeable, non-fluorescent DHR 123 dye into green fluorescent rhodamine upon stimulation with PMA (Phorbol 12-myristate 13-acetate) by flow cytometry. The cell gating strategy in each condition is shown in Fig EV1A, 1B. Our data showed that after stimulation with PMA, the median fluorescence intensity was higher under each of the induced conditions compared to uninduced (stimulated) HL-60 cells (Fig 1C). Similarly, cells induced with DMSO (p=0.03) and DMSO+D3 (p=0.01) exhibited significantly higher neutrophil oxidative index values, compared to the uninduced HL-60 cells (Fig 1D).

### DMSO+D3-induced HL-60 cells express surface antigens and additional markers characteristic of immature neutrophils

Flow cytometric analysis was performed to assess the expression of CD11b (myeloid lineage marker), CD16 (mature granulocyte marker) and CD14 (marker of immature granulocytes, monocytes and M2 macrophages). The gating strategy of cells is shown in Fig EV1C-E. Fig 2 shows that uninduced HL-60 cells predominantly exhibited a CD11b^-^CD14^-^ (99.7%) or CD11b^-^CD16^-^ (99.7%) phenotype, consistent with their promyelocytic state. Upon induction with various inducers, cells showed increased CD11b expression, indicating differentiation towards myeloid lineage. HL-60 cells induced with a combination of DMSO and D3, displayed a higher percentage (87%) of CD11b^+^CD14^+^cells (Fig 2A), indicative of immature neutrophils, and a low percentage (0.9%) of CD11b^+^CD16^+^ cells (Fig 2B), indicative of mature neutrophils. CD11b^+^CD14^+^cells were also observed in HL-60 cells induced with D3 alone (6.7%) (Fig 2A). Notably, CD16 expression was not detected under any condition (Fig 2B).

**Figure 2.**
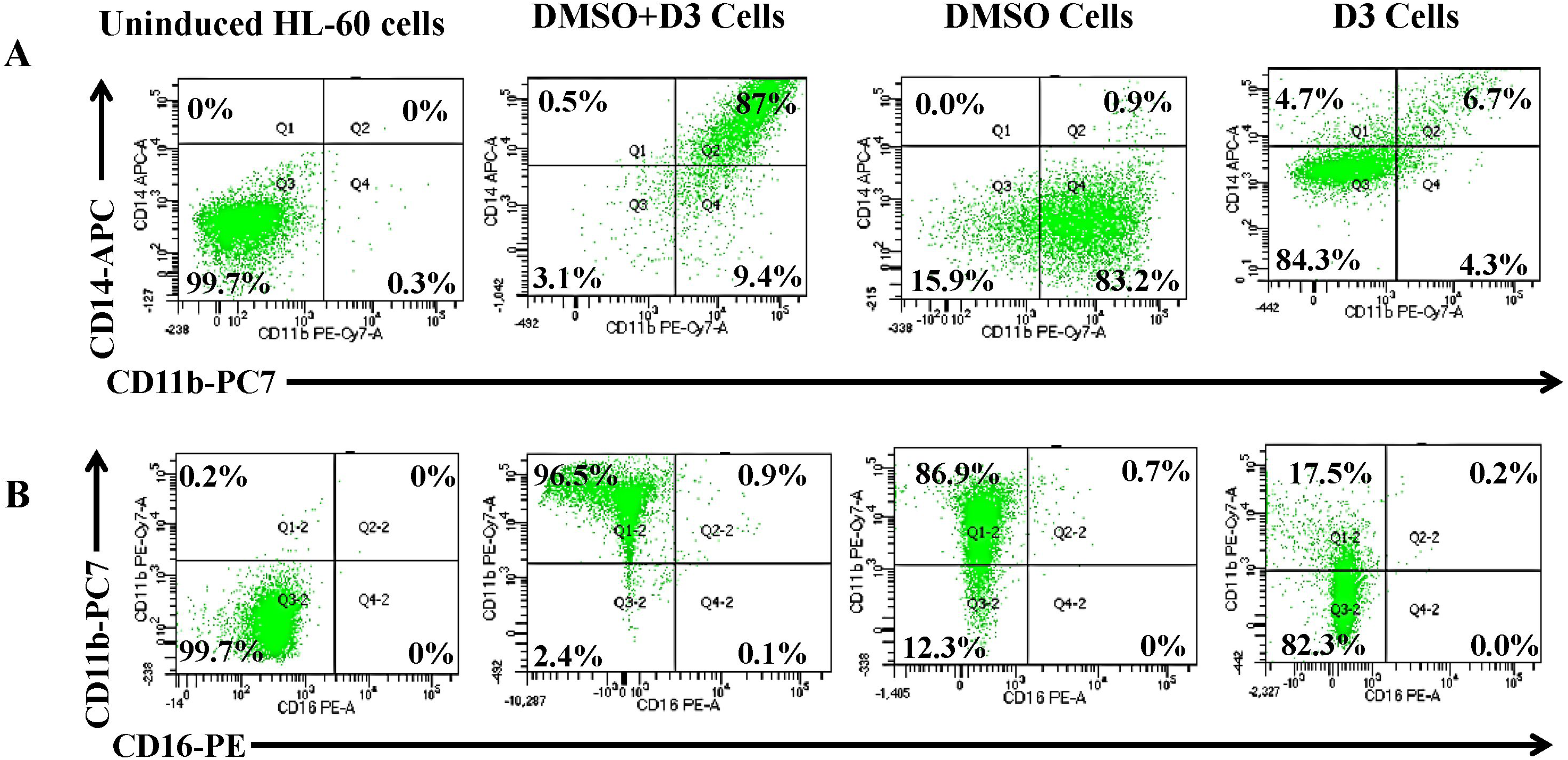
Expression of surface immune cell markers in HL-60 cells, upon induction with different reagents. Representative flow cytometric analysis, showing the expression of (A) CD11b+CD14+ and (B) CD11b+CD16+ cell surface markers in uninduced HL-60 cells, or upon induction with DMSO+D3, DMSO alone or D3 alone for 4 days. Data are representative of three independent experiments (n=3).

In order to determine whether prolonged exposure to DMSO+D3 could drive HL-60 differentiation towards mature neutrophils, the expression of these markers was also analyzed on days 6 and 8 in uninduced HL-60 cells or upon induction with DMSO and DMSO+D3, using the gating scheme shown in Fig EV2. Cells induced with DMSO+D3 maintained an increased percentage of CD11b^+^CD14^+^cells on day 6 (87%) and day 8 (89.2%), as seen on day 4 (80.6%) (Fig EV3A). In contrast, the population expressing CD11b^+^CD16^+^ remained low across all the time points, i.e., 0.1% at day 4, 1.6% at day 6 and 6.8% at day 8, indicative of maintenance of immature neutrophil-like phenotype over time, rather than differentiation into mature neutrophils (Fig EV3B).

Further, expression of these markers was also assessed at transcript level and compared with uninduced HL-60 cells (Fig 3). It was observed that expression of CD11b was significantly increased under all culture conditions (DMSO: p=0.004; DMSO+ D3: p=0.03; D3: p=0.01). Conversely, CD16 expression was significantly downregulated upon induction of HL-60 cells with DMSO (p=0.0008); DMSO+ D3 (p=0.001); and D3 (p=0.006). CD14 expression was found to be increased in cells induced with DMSO+D3 (p=0.03), while it was significantly decreased in HL-60 cells induced with DMSO (p=0.01) (Fig 3), in line with the flow cytometry data. Moreover, the increase in CD14 expression in DMSO+D3-induced cells was significantly higher as compared to cells induced with either DMSO alone (p=0.02) or D3 alone (p=0.04).

**Figure 3.**
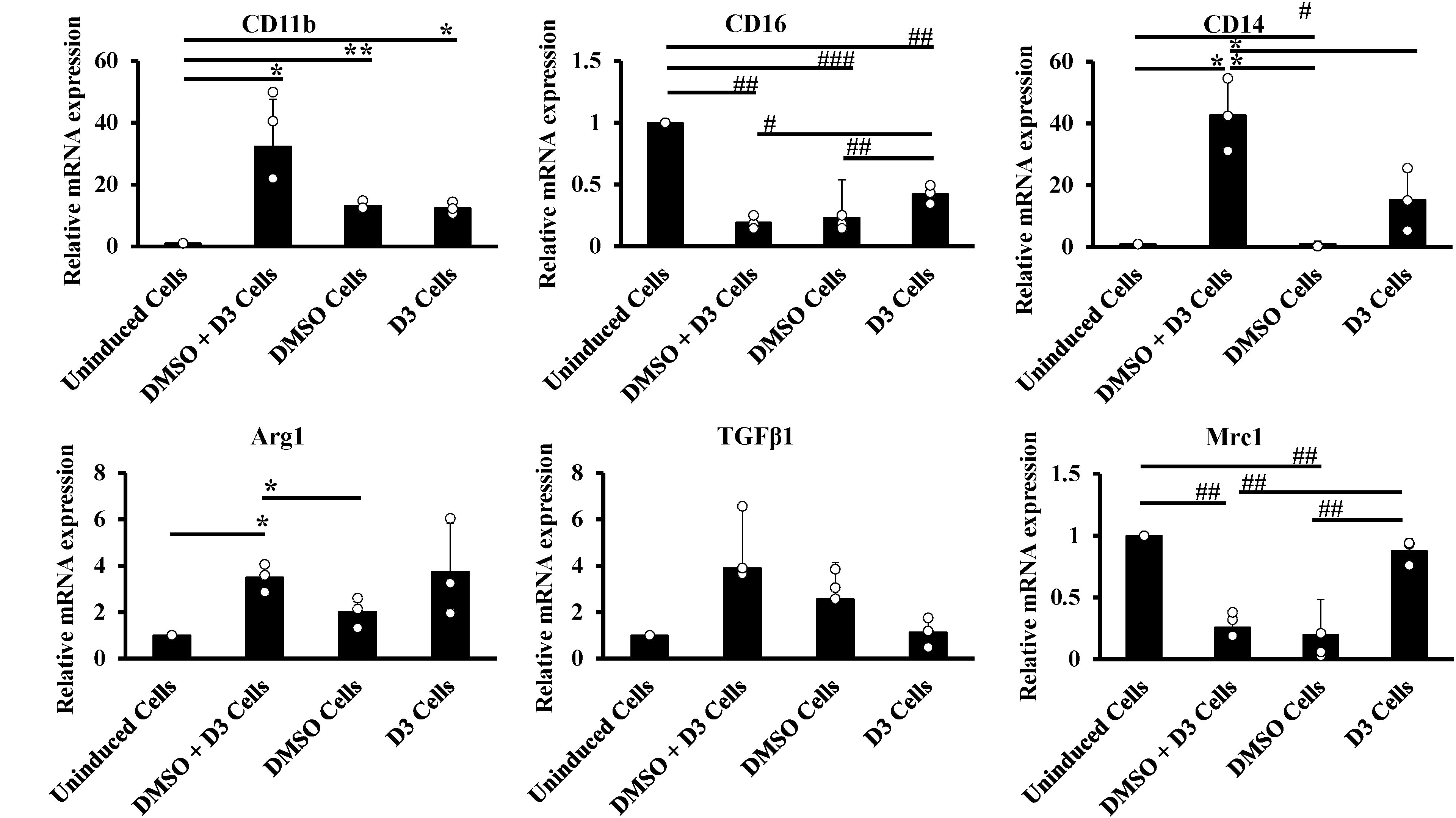
Transcript expression of immune cell-related markers in HL-60 cells, upon induction with different reagents. Relative mRNA expression levels of CD11b, CD16, CD14, arginase 1 (ARG1), mannose receptor 1 (MRC1) and TGFβ1 were analysed by quantitative real-time PCR (qRT-PCR) in uninduced HL-60 cells or upon induction with DMSO + D3, DMSO alone or D3 alone for 4 days; expression of genes in each case was normalized with respect to the uninduced HL-60 cells. Data are represented as mean ± standard deviation from three independent experiments (n=3). Statistical significance was calculated using Student’s t-test; *P* ≤0.05 was considered significant [\**P* ≤ 0.05, \*\**P* < 0.01, \*\*\**P* < 0.001; symbols (*) and (#) indicate increased and decreased expression respectively].

Transcript expression of other markers, such as arginase 1 (expressed by M2 macrophages and immature neutrophils), TGFβ1 (associated with M2 macrophages), and MRC1 (expressed by M2 macrophages), were also assessed. DMSO+ D3-induced HL-60 cells showed significantly elevated expression of arginase 1 (p=0.02) as compared to uninduced cells; this expression in DMSO+D3-induced cells was also significantly increased as compared to DMSO-induced HL-60 cells (p=0.04). TGFβ1 expression was highest in DMSO+D3-induced HL-60 cells, followed by other conditions, although the changes were not statistically significant (ns; p=0.19). In contrast, MRC1 expression was decreased under all culture conditions (DMSO: p=0.003; DMSO+ D3: p=0.001; D3: ns-p=0.16), as compared to induced cells. Additionally, DMSO (p=0.001) and DMSO+D3-induced HL-60 cells (p=0.002) showed decreased expression of MRC1 as compared to D3-induced HL-60 cells (Fig 3).

### Proteome analyses of DMSO+D3-induced HL-60 cells reveal signature of immature neutrophil-like cells along with proteins associated with wound healing

We carried out a label free quantitative proteome analysis of uninduced HL-60 cells, and HL-60 cells induced with DMSO+D3. The principal component analysis (PCA) plot showed a distinct separation between the uninduced and induced cell populations (Fig 4A), forming clearly defined clusters. A total of 2,008 proteins were detected, of which 467 were upregulated (fold change ≥ 1.5), 1292 were downregulated (fold change <1.5), 103 were uniquely expressed in DMSO+D3-induced HL-60 cells, 28 were uniquely expressed in uninduced HL-60 cells, and 118 proteins were undefined. Reactome pathway analysis indicated that DMSO+D3-induced HL-60 cells expressed an increased percentage of proteins associated with the innate immune system, metabolism, neutrophil degranulation, signaling by Rho GTPase, and Fc-gamma receptor mediated phagocytosis (Fig 4B). These cells showed increased expression of proteins associated with immature neutrophils (PTPRC, ITGAM, CD14), ROS production (NCF2, NCF4, CTSB, G6PD, SOD2, CTSD, HCK, PRKCB, CYBB, RAC1, RAC2 and PRDX5), and neutrophil activation and migration (S100A8, S100A9, ANXA5) as listed in Table 1 and Fig 4C.

**Figure 4.**
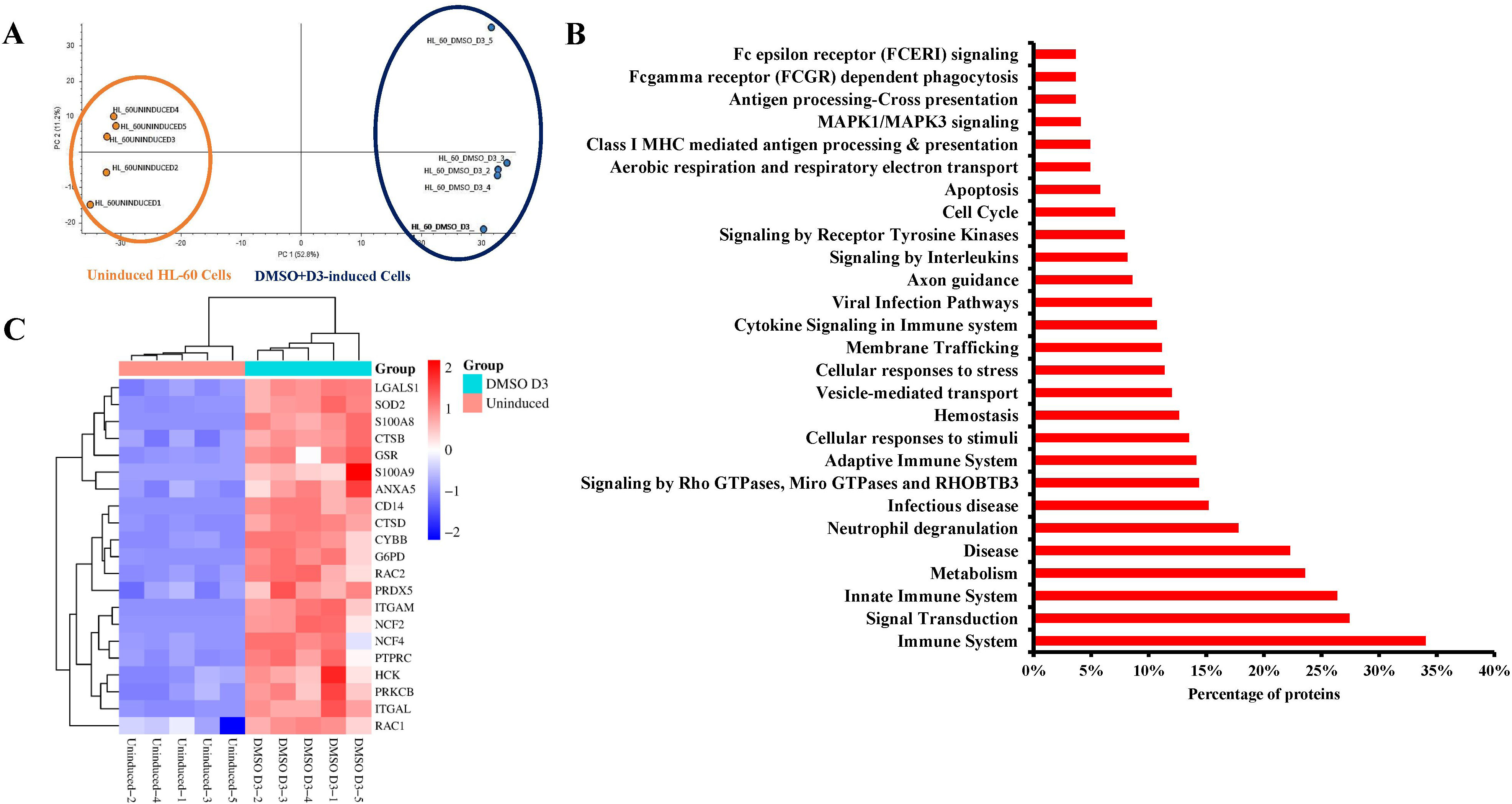
Mass spectrometry-based proteomic comparison of HL-60 cells, under uninduced condition, or upon induction with DMSO+D3. (A) Principal component analysis (PCA) plot demonstrating the distinct separation of proteomic expression patterns in both the culture conditions, with uninduced HL-60 replicates represented in orange, and DMSO+D3-induced samples in blue colour. (B) Top differentially expressed Reactome pathways, generated based on the increased percentage of proteins and (C) heat map showing proteins related to neutrophil biology in DMSO + D3-induced HL-60 cells compared to uninduced cells. Proteomic analysis was performed using five biological replicates per condition (n=5). Differential protein expression was assessed by calculating fold changes based on the average normalized abundances of HL-60 cells induced with DMSO+D3, relative to uninduced HL-60 cells. Proteins with fold change ≥1.5, along with a false discovery rate–adjusted P-value less than 0.05 (*P* <0.05), were considered to be differentially expressed and considered for analysis.

**Table 1:**
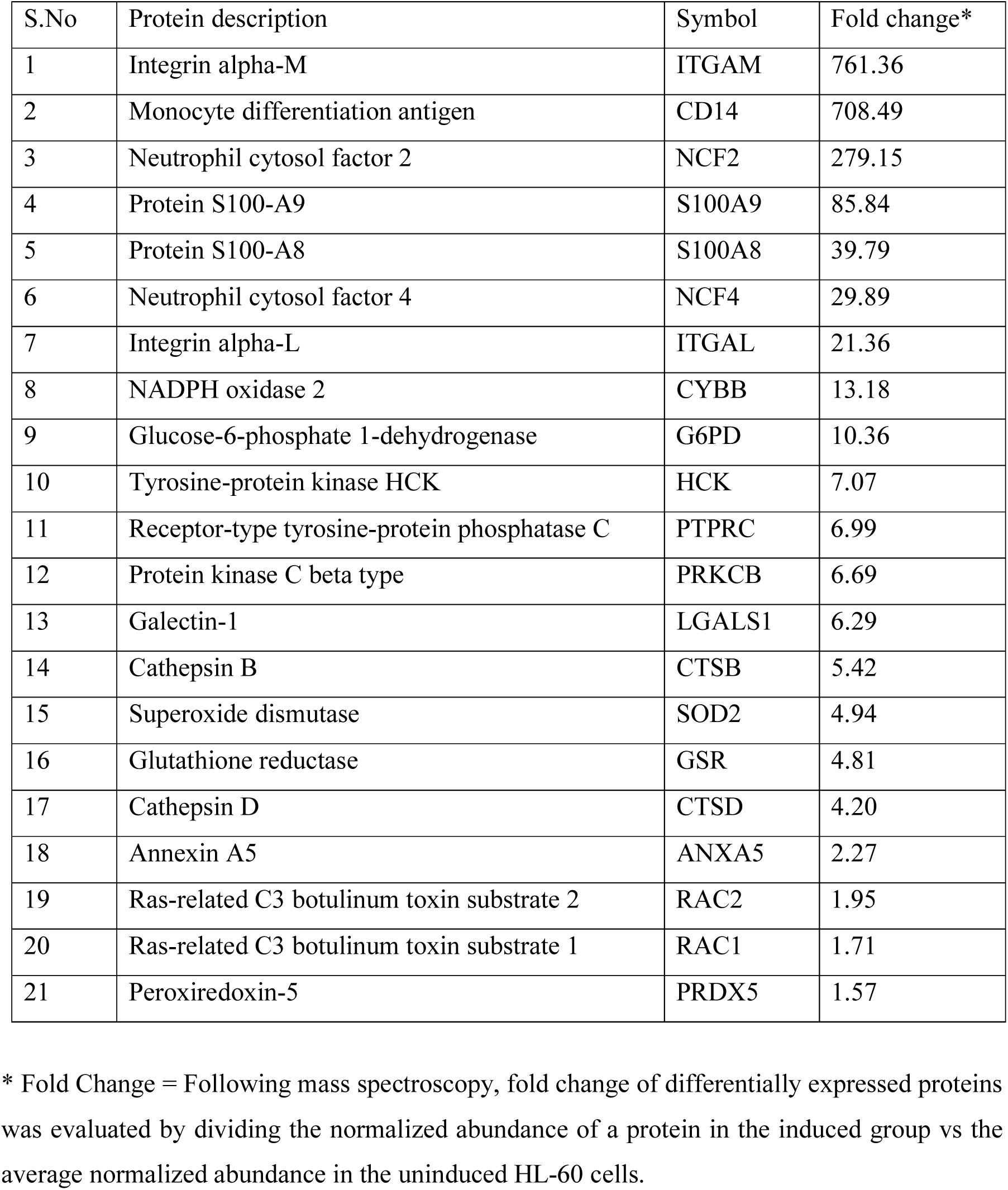
Fold change of differentially expressed proteins in DMSO+D3-induced HL-60 cells versus uninduced HL-60 cells, associated with various functions related to neutrophil biology.

| S.No | Protein description | Symbol | Fold change* |
| --- | --- | --- | --- |
| 1 | Integrin alpha-M | ITGAM | 761.36 |
| 2 | Monocyte differentiation antigen | CD14 | 708.49 |
| 3 | Neutrophil cytosol factor 2 | NCF2 | 279.15 |
| 4 | Protein S100-A9 | S100A9 | 85.84 |
| 5 | Protein S100-A8 | S100A8 | 39.79 |
| 6 | Neutrophil cytosol factor 4 | NCF4 | 29.89 |
| 7 | Integrin alpha-L | ITGAL | 21.36 |
| 8 | NADPH oxidase 2 | CYBB | 13.18 |
| 9 | Glucose-6-phosphate 1-dehydrogenase | G6PD | 10.36 |
| 10 | Tyrosine-protein kinase HCK | HCK | 7.07 |
| 11 | Receptor-type tyrosine-protein phosphatase C | PTPRC | 6.99 |
| 12 | Protein kinase C beta type | PRKCB | 6.69 |
| 13 | Galectin-1 | LGALS1 | 6.29 |
| 14 | Cathepsin B | CTSB | 5.42 |
| 15 | Superoxide dismutase | SOD2 | 4.94 |
| 16 | Glutathione reductase | GSR | 4.81 |
| 17 | Cathepsin D | CTSD | 4.20 |
| 18 | Annexin A5 | ANXA5 | 2.27 |
| 19 | Ras-related C3 botulinum toxin substrate 2 | RAC2 | 1.95 |
| 20 | Ras-related C3 botulinum toxin substrate 1 | RAC1 | 1.71 |
| 21 | Peroxiredoxin-5 | PRDX5 | 1.57 |

\* Fold Change = Following mass spectroscopy, fold change of differentially expressed proteins was evaluated by dividing the normalized abundance of a protein in the induced group vs the average normalized abundance in the uninduced HL-60 cells.

### DMSO+D3-induced HL-60 cells promote wound closure in human corneal epithelial cells (HCEC)

Given the importance of cell migration in re-epithelialization, a scratch-wound assay using a transwell co-culture system was used to examine the effect of these immature neutrophil-like cells on corneal epithelial wound healing (Kacham et al, 2021), through measurement of wound closure in cultured monolayer of human corneal epithelial cells (HCEC). It was observed that relative to the serum-free media control, HL-60 cells induced with DMSO+D3 (p=0.004) or D3 alone (p=0.04) resulted in a significant increase in percentage wound closure of the HCEC cells. Notably, DMSO+D3-induced immature neutrophil-like cells exhibited significantly increased percentage of wound closure compared to uninduced HL-60 cells (p=0.01) as well as when compared to HL-60 cells induced with DMSO alone (p=0.04) or D3 alone (p=0.01) (Fig 5 A, B).

**Figure 5.**
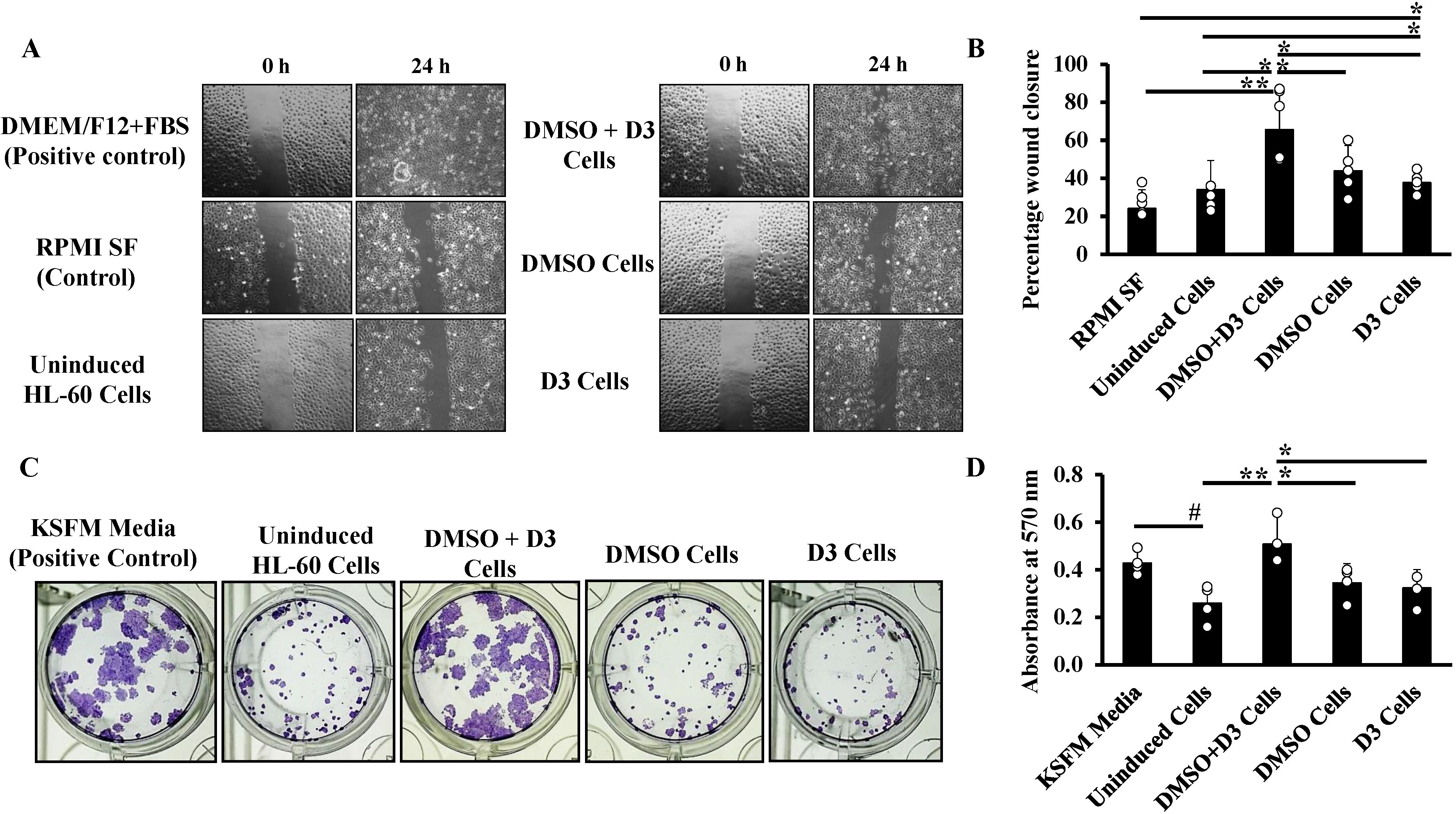
DMSO+D3-induced HL-60 cells promote corneal epithelial wound closure and limbal epithelial cell colony formation. (A) Representative phase-contrast images of wound in scratch assay at 0 h and 24 h captured by light microscope at 10X magnification (scale bar: 400 µm) and (B) quantification of percentage wound closure of HCEC (added in the bottom chamber of transwell inserts) after co-culture with HL-60 uninduced cells and HL-60 cells induced with various reagents (added in top chamber) for 24 h, compared with RPMI SF media control. Cells cultured in the presence of FBS served as positive control. (C) Representative images of HCLE colonies formed, and (D) corresponding quantitative measurement of crystal violet uptake at 570 nm in HCLE cells (added in the bottom chamber of transwell inserts) after co-culture with HL-60 cells, uninduced and induced with various reagents (added in top chamber) for seven days. Data in (B, D) are represented as mean ± standard deviation from four independent experiments (n=4). Statistical significance was calculated using Student’s t-test; *P* ≤0.05 was considered significant [\**P* ≤ 0.05, \*\**P* < 0.01; symbols (*) and (#) indicate increased and decreased expression respectively].

### DMSO+D3-induced immature neutrophil-like cells promote colony formation of human corneal limbal epithelial (HCLE) cells

Limbal epithelial stem cells (LESCs) are a population of stem cells, characterized by a slow cell cycle, asymmetric division, low differentiation rate, and high proliferative and self-renewal capability. These cells function to protect and repair the corneal integrity by continuous renewal and regeneration of the corneal epithelium. We tested the ability of DMSO+D3-induced HL-60 cells to promote colony formation of human corneal limbal epithelial cell line (HCLE), used here as an *in vitro* mimic for LESCs in a transwell based indirect co-culture system (Nam et al, 2017). It was observed that HCLE cells co-cultured with DMSO+D3-induced HL-60 cells showed increased colony formation compared to other (induced) conditions and exhibited colonies comparable to, or exceeding those observed in KSFM medium (positive control) (Fig 5C). Quantification by crystal violet staining (Kumar et al, 2018; An et al, 2021) further complemented these observations, indicating increased absorbance in DMSO+D3-induced HL-60 cells under the co-culture condition, which correlates with enhanced colony formation, as compared to uninduced HL-60 cells (p=0.006), and HL-60 cells induced with DMSO (p=0.04) or D3 (p=0.02) (Fig 5D).

### DMSO+D3-induced immature neutrophil-like cells are effective in healing of alkali-injured goat corneas (*ex vivo*)

DMSO+D3-induced immature neutrophil-like cells were applied to alkali (sodium hydroxide; NaOH)-injured goat corneas, cultured *ex vivo* (Shukla et al, 2024); corneal wound healing and tissue restoration was assessed over a period of 7 days, by examining epithelial healing using fluorescein staining, histological examination and immunofluorescence evaluation of the corneal epithelial marker KRT12 and the fibrotic marker, α-SMA.

In contrast to uninjured goat corneas, alkali-injured goat corneas showed intense fluorescein staining indicating prominent epithelial defects caused by alkali burn at day 0. Alkali-injured goat corneas treated with RPMI SF media served as controls, showing persistent epithelial defect at day 7. Notably, treatment of alkali-injured goat corneas with DMSO+D3-induced HL-60 cells (Fig 6A) showed absence of fluorescein staining by day 7, indicating complete epithelial healing, thereby suggesting that these cells promote effective epithelial healing, and maintain epithelial integrity following alkali injury. Additionally, uninjured goat corneas showed normal histology with stratified epithelium, intact Bowman’s layer, organized stromal collagen fibrils (populated with keratocytes), defined Descemet’s membrane and endothelium. In contrast, alkali-injured goat corneas, treated with media, showed epithelial loss and hyalinization of stromal layer, characterized by tissue degeneration into a translucent, glass-like appearance. Remarkably, treatment of alkali-injured goat corneas with DMSO+D3-induced immature neutrophil-like cells showed restoration of epithelial layer and stromal architecture, indicating enhanced tissue repair (Fig 6B). In line with these results, uninjured corneas showed presence of a continuous epithelium lining, expressing the epithelial marker, KRT12. Similarly, alkali-injured goat corneas treated with DMSO+D3-induced HL-60 cells, showed formation of epithelium lining, with expression of KRT12 as compared to media control (Fig 6C). Immunofluorescence expression of α-SMA, a marker of myofibroblast activation and fibrosis was performed to assess the extent of fibrotic remodelling. Uninjured corneas showed minimal to no expression of α-SMA, whereas alkali-injured goat corneas treated with RPMI SF media exhibited elevated expression, indicative of fibrotic changes. Compared to media control, alkali-injured corneas, treated with DMSO+D3-induced HL-60 cells showed reduced α-SMA expression, suggesting reduction in fibrosis and improved stromal remodelling (Fig 6D).

**Figure 6.**
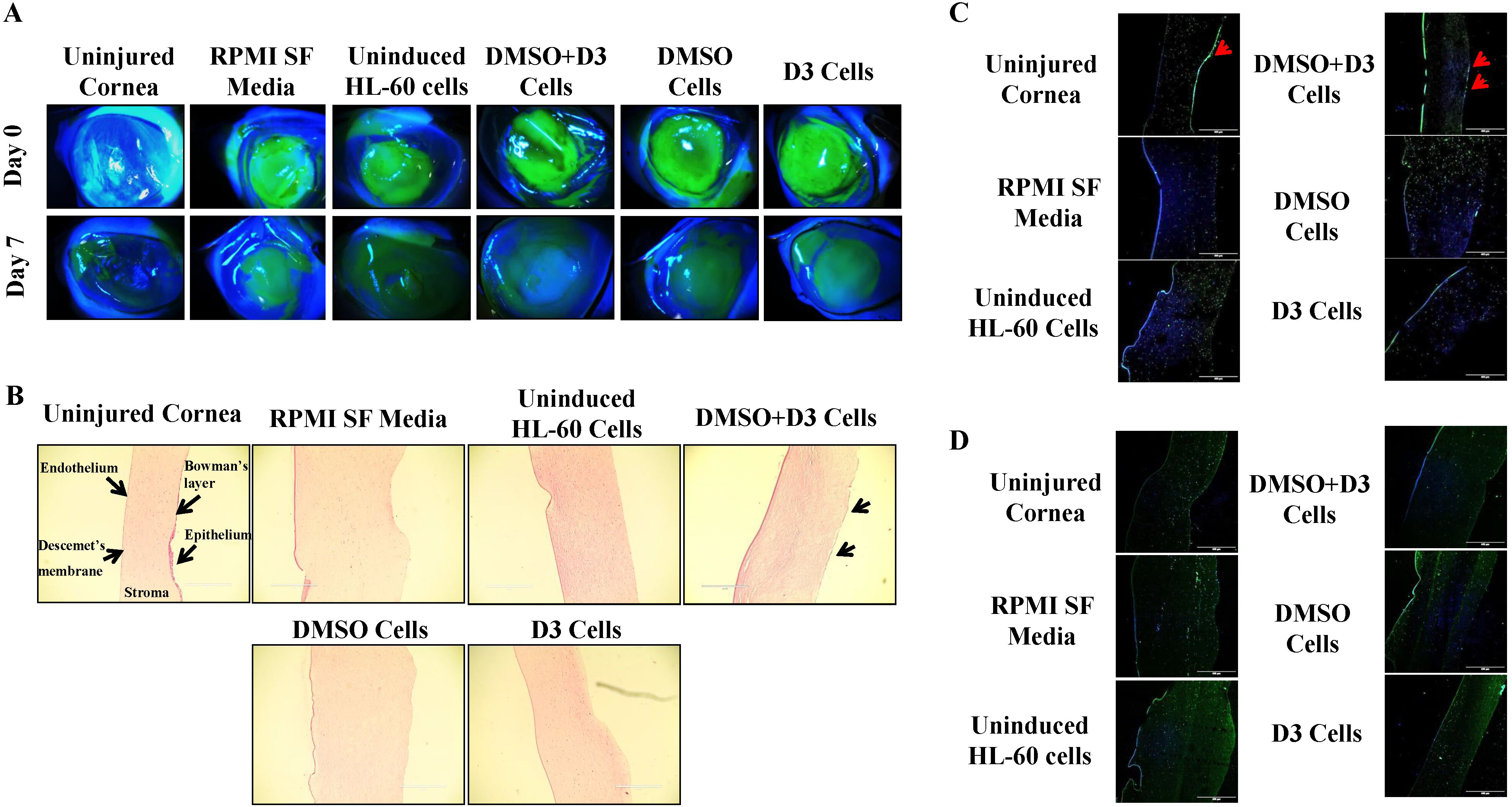
Promotion of corneal repair in alkali-injured goat corneal organ cultures following treatment with DMSO+D3-induced HL-60 cells. Goat corneas were injured with alkali, followed by administration of vehicle alone (RPMI serum free media) or HL-60 cells, either uninduced or induced with DMSO + D3, DMSO alone or D3 alone. (A) Representative slit-lamp images acquired at 10 X magnification (scale bar: 400 µm) on day 0 and day 7; (B) H&E images following termination of the experiment on day 7, acquired by bright-field microscopy at 10 X magnification (scale bar: 400 µm). Arrows in test groups point towards formation of corneal epithelium; Immunofluorescence analysis of (C) epithelial marker KRT12, and (D) fibrotic marker, α-SMA. Images were acquired by confocal microscopy at 10 X magnification (scale bar: 400 µm). Merged images are shown; DAPI: nuclear staining, Alexa fluor 488 for KRT12 marker as well as α-SMA markers. Data are representative of two independent experiments (A, B) and one experiment with n=3 corneas (C, D).

### DMSO+D3-induced immature neutrophil-like cells promote re-epithelialization and reduce post-injury fibrosis in corneal burn mice model

We examined the efficacy of DMSO+D3-induced immature neutrophil-like cells in healing of corneas in the alkali burn murine model established in C57BL/6 female mice (Jhanji et al, 2022; Santra et al, 2024).

Corneal epithelial defect, opacity and neovascularization scoring was carried out by an ophthalmologist (in a blinded fashion) at day 0, 7 and day 14, post-injury. Following the alkali burn, the corneal epithelial defect and opacity were scored on a scale of 0 to 4. Post-injury on day 0, the mice in both the groups (vehicle-treated and DMSO+ D3 cells-treated) had comparable injury to both the layers of cornea, i.e., the corneal epithelium and stroma. The average epithelial defect score was 3, representing 50% to 75% epithelial defect, while stromal opacity showed an average score of 2, depicting slightly opaque stroma, with iris and pupil still detectable (Fig 7A; Fig EV4A, 4B), indicating uniformity of the injury induced. Complete re-epithelialization was observed in both the groups by day 14, as observed by absence of fluorescein staining and an average epithelial defect score of 0 (Fig 7A; Fig EV4A). Given that the epithelial layer restoration occurred in both the groups, indicating effectiveness towards epithelial wound healing, differences were assessed in opacity recovery. Corneal opacity was found to be increased in both the groups at day 7; however, it was lower in DMSO+D3-treated group (ns: p=0.53) compared to vehicle group. Notably, by day 14, the opacity reduced significantly in the group treated with cells compared to vehicle group (p=0.03). In contrast, on day 14, the opacity observed in vehicle group was increased, when compared to vehicle (p=0. 0.0006) group and the treatment group (p=0.001) on day 0 (Fig 7A and Fig EV4B). Further, corneal neovascularization was scored on a scale of 0-3. The formation of neo-vessels was observed in both the groups, with reduction on day 14 in DMSO+D3 cells-treated group compared to vehicle group (ns: p=0.31), signifying a protective effect of the cells against angiogenesis (Fig EV4C). Histological analysis performed on day 14-post injury, showed that while the uninjured mouse corneas displayed intact corneal integrity, the alkali injured mouse corneas in vehicle group, showed pronounced pathological changes, such as corneal epithelial thinning, degenerative changes in the stroma with disorganized collagen fibrils, moderate edema characterized by thickening of cornea and presence of inflammatory cells (Fig 7B). On the other hand, injured corneas treated with DMSO+D3-induced HL-60 cells, showed significant restoration of tissue architecture, with mild corneal epithelial thinning, restoration of damaged epithelium with formation of epithelial layers, organized stromal collagen, mild edema, and reduced inflammatory cell infiltration (Fig 7B). These findings are in correlation with reduced opacity as well as neo-vessel formation observed during clinical examination.

**Figure 7.**
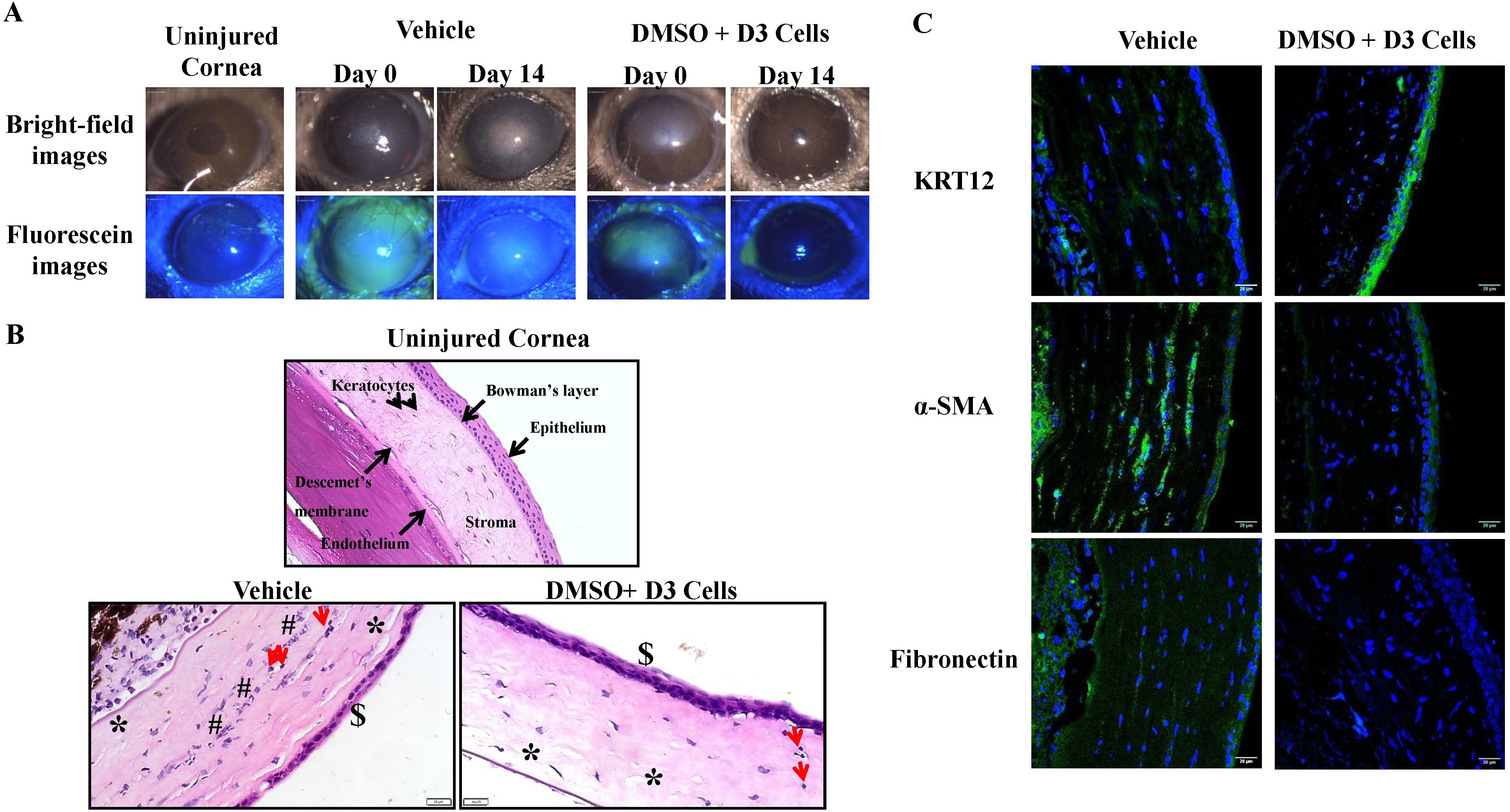
DMSO+D3-induced HL-60 cells enhance corneal wound healing *in vivo*. Mice corneas were alkali-injured and treated with either vehicle alone (fibrin glue only) or DMSO+D3-induced HL-60 cells. (A) Representative bright-field images showing corneal opacity (top row) and fluorescein-stained epithelial defects (bottom row), captured under diffuse and cobalt blue light illumination, respectively, captured at 40 X magnification by slit lamp microscope (scale bar: 100 µm); (B) Representative H&E images of uninjured and alkali-injured mouse corneas treated with either vehicle or DMSO+ D3-induced HL-60 cells on day 14, post-injury. The symbols indicate corneal epithelial thinning ($), degenerative stromal changes (#), edema (*) and infiltration of immune cells (red arrows). Images were acquired using bright-field microscope at 40 X magnification (scale bar: 100 µm); (C) assessment of expression of corneal epithelial marker, KRT12, and fibrotic markers, α-SMA and fibronectin, in alkali-injured mouse corneas, following the respective treatments on day 14, post injury. Merged images are shown; DAPI: nuclear staining, Alexa fluor 488 for KRT12, α-SMA and fibronectin markers. Images were acquired by confocal microscopy at 60 X magnification (scale bar: 20 µm). (B, C) Data is represented from one experiment; n=3 eyes in the vehicle group and n=4 eyes in the cells-treated group.

In line with the findings of clinical and histological examination, indicating the restoration of epithelium in alkali-injured mouse corneas treated with DMSO+D3-induced HL-60 cells, immunofluorescence analysis showed increased expression of KRT12 in alkali injured corneas treated with cells (Fig 7C), confirming the regeneration of corneal epithelium. Compared to vehicle-treated controls, corneas treated with DMSO+D3-induced HL-60 cells showed reduced α-SMA and fibronectin expression (Fig 7C), demonstrating attenuation of stromal fibrosis in this group.

### Evidence of presence of reparative macrophage phenotype in alkali injured corneas treated with DMSO+D3-induced immature neutrophil-like cells

After an injury to the cornea, immune cells such as neutrophils and macrophages enter the cornea, resulting in a cascade of inflammatory reactions and delayed wound repair (Wilson et al, 2001). Modulation of immune cell recruitment, specifically inhibition of neutrophil infiltration (Hertsenberg et al, 2017; Santra et al, 2024), and promotion of phenotypic transition of macrophages towards an alternatively activated, anti-inflammatory M2 macrophages instead of pro-inflammatory M1 macrophages is important for resolution of corneal pathology (He et al, 2023). Consequently, transcript analysis was carried out to check the expression of neutrophils (Ly6G), macrophages (F4/80), M1 macrophages (CD86) and M2 macrophages (CD163) in vehicle-treated and cells-treated alkali-injured mouse corneas.

It was observed that mouse corneas administered with DMSO+D3-induced HL-60 cells showed significant suppression of neutrophil infiltration, as indicated by reduced Ly6G levels (p= 0.00003) (Fig 8A). On the other hand, an increase in macrophage cell population was observed (F4/80; p=0.006) (Fig 8B). However, although increase in both M1 (CD86; p=0.03) (Fig 8C) and M2 (p=0.0002) (Fig 8D) macrophage populations was observed, the M2 macrophage population predominated over the M1 population (p=0.03) (Figure 8E). This transition towards an alternative, reparative macrophage phenotype in mice corneas treated with DMSO+D3-induced HL-60 cells, explains the decrease in inflammation, fibrosis and enhanced corneal repair.

**Figure 8.**
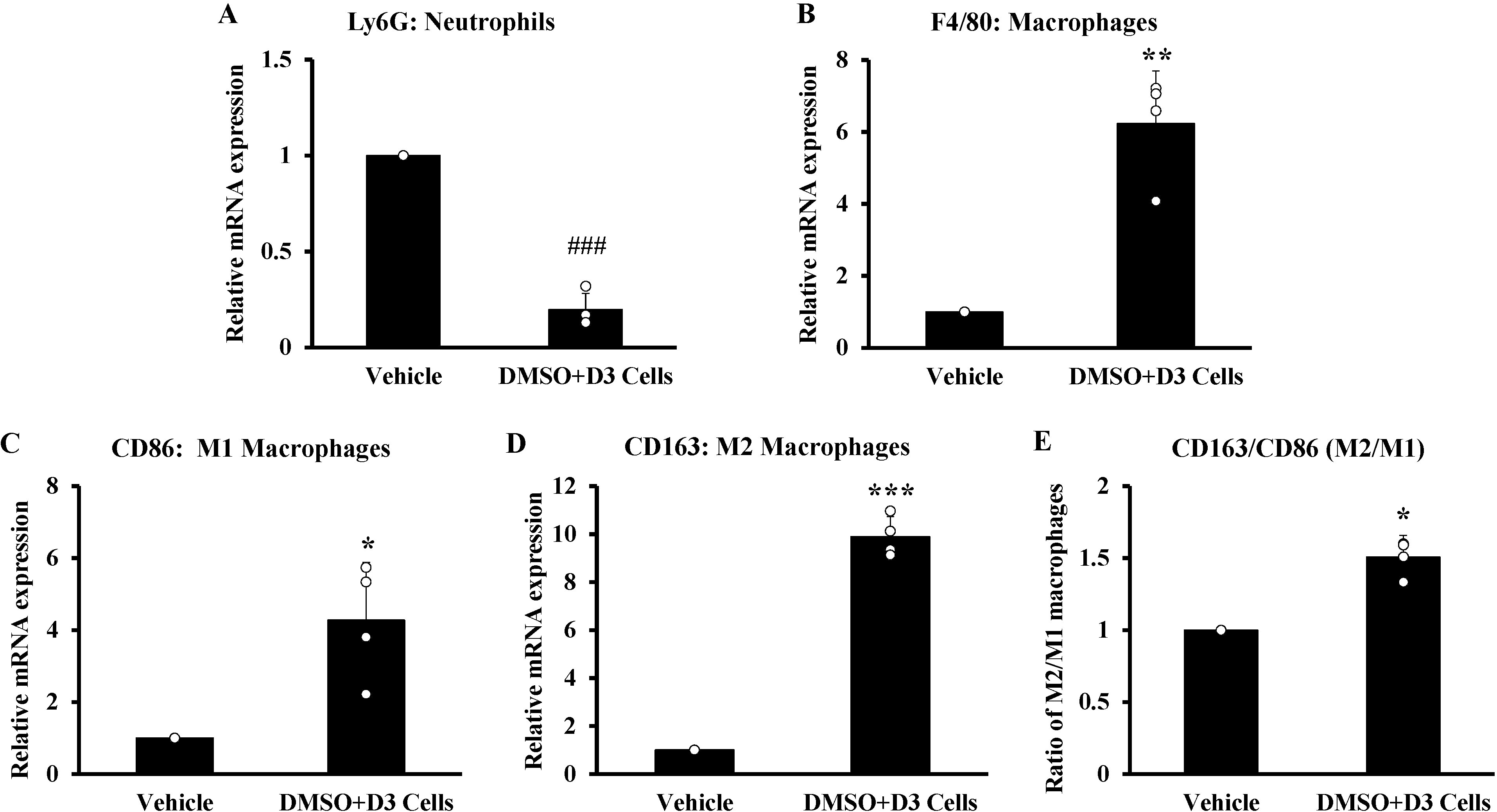
Immunomodulatory effects of DMSO+D3-induced HL-60 cells in alkali-injured mouse corneas. Relative mRNA expression of (A) the neutrophil marker: Ly6G, (B) macrophage marker: F4/80, (C) M1 macrophage marker: CD86, (D) M2 macrophage marker: CD163 and (E) the M2/M1 macrophage ratio in alkali-injured mouse corneas, treated with either vehicle alone (fibrin glue only) or DMSO+D3-induced HL-60 cells on day 14 post-injury, as compared with vehicle-treated corneas (control). Data are represented as mean ± standard deviation from one experiment, with n=3 eyes in the vehicle group and n=6 eyes in the cells-treated group. Statistical significance was calculated using Student’s t-test; *P* ≤0.05 was considered significant [\**P* ≤ 0.05, \*\**P* < 0.01, \*\*\**P* < 0.001; symbols (*) and (#) indicate increased and decreased expression respectively].

## Discussion

In this study, we have developed an easy and reproducible protocol for generating high numbers of immature neutrophil-like cells from HL-60 cells by using a combination of DMSO and 1α,25-dihydroxyvitamin D3. A comprehensive analysis was carried out for validating the success of the differentiation status and functional identity of these cells. Further, the repair and regenerative capabilities of these cells was assessed in the context of corneal wound healing.

Morphologically, uninduced HL-60 cells exhibited promyelocytic features, consistent with their origin, displaying a uniform shape and a large nuclear-to-cytoplasmic ratio even after four days of culture, thereby confirming the absence of differentiation. In contrast, HL-60 cells induced with DMSO+D3 showed features consistent with differentiation towards neutrophils, including decreased cell size, loss of uniformity, presence of membrane extensions (Gee et al, 2012), and band-shaped nuclei with a reduced nuclear-to-cytoplasmic ratio among all the tested conditions (He et al, 2025; Gysemans et al, 2025).

Neutrophils are a highly heterogenous population, exhibiting variability in marker expression between immature and mature phenotypes. Therefore, no single specific marker can delineate a particular neutrophil phenotype. Human mature neutrophils with segmented nuclei and high density, display higher expression of CD10, CD11b, CD16 and CD62L (Gysemans et al, 2025), whereas low density neutrophils (LDNs) exist as banded, immature or segmented activated populations. Immature banded LDNs show decreased expression of CD16 and exhibit increased expression of CD66b and CD33. However, activated LDNs have increased expression of CD64 and CD54, while retaining CD16 expression (Blanco-Camarillo et al, 2021; Gysemans et al, 2025). Granulocyte-myeloid derived suppressor cells (G-MDSCs) express CD11b^+^, CD15^+^, CD16^+^, CD14^−^ and HLA-DR^−^ markers (Cassetta et al, 2019; Blanco-Camarillo et al, 2021). Neutrophils can also be polarized into pro-inflammatory N1 neutrophils, expressing CD11b, CD66b and CD64, or anti-inflammatory N2 neutrophils, expressing CD163, CD16 and CD206 (He et al, 2025). Moreover, a subset of neutrophils with immunoregulatory or reparative roles have been characterized in mice, possessing features of immature neutrophils with band-shaped nuclei and expression of Ly6G^low^CD14^+^ along with M2-like macrophages-associated markers such as Arg1, Mrc1, Tgfb1, Igf1, and Ccl5 (Sas et al, 2020; Jerome et al, 2022).

Considering these parameters, we designed a panel of markers corresponding to mature and immature neutrophils and assessed their expression at both the protein (CD11b, CD14 and CD16) and transcript (CD11b, CD14, CD16, arginase 1, TGFβ1, and MRC1) levels to define the generation of immature neutrophil-like cells. Flow cytometric as well as transcript expression data confirmed the promyelocytic features of uninduced HL-60 cells, which were negative for all assessed markers. In contrast, HL-60 cells induced with DMSO+D3 showed increased expression of CD11b^+^CD14^+^, indicating an immature phenotype, and decreased expression of CD11b^+^CD16^+^, indicative of a mature phenotype.

Proteomic analysis further confirmed the immature neutrophil-like phenotype of these cells, revealing upregulation of several proteins associated with neutrophil biology. The increased expression of adhesion proteins, such as integrin alpha-M (CD11b) and integrin alpha-L (CD11a), indicates the potential of these cells for adhesion and migration (Chidlow et al, 2010). Furthermore, calcium-binding proteins S100A8 and S100A9, which form the calprotectin complex, found in cytoplasmic compartment of neutrophils, functioning as amplifiers of neutrophil activation were found to be upregulated (Sprenkeler et al, 2022). Proteins involved in the assembly of the NADPH oxidase complex-neutrophil cytosol factor 2 (p67^phox^), neutrophil cytosol factor 4 (p40^phox^), NADPH oxidase 2 (NOX2/gp91^phox^) and small GTPases, RAC1 and RAC2 were also upregulated, suggesting the formation of a functional oxidative burst machinery (Paclet et al, 2022). Additionally, glucose-6-phosphate dehydrogenase (G6PD), the rate-limiting enzyme of the pentose phosphate pathway that provides NADPH for ROS production was detected (Paclet et al, 2022). Functionally, these cells exhibited enhanced ROS production upon PMA stimulation, confirming their differentiation towards the myeloid lineage, specifically towards granulocytes.

Moreover, annexin a5, an anti-inflammatory molecule (Tschirhart et al, 2023; Kang et al, 2024) and galectin-1, a protein involved in various neutrophil effector functions (Robinson et al, 2019) were also upregulated. Importantly, CD14 was found to be highly upregulated, indicating the generation of immature neutrophil-like cells with active ROS production and expression of key proteins related to neutrophil biology.

It is noteworthy that DMSO+D3-induced HL-60 cells are effective in promoting colony formation of LESCs. Damage to LESCs or disruption of their niche results in a pathological condition, called limbal stem cell deficiency (LSCD), wherein the barrier between corneal and conjunctival epithelium is disrupted, resulting in the invasion of the latter into the cornea and failure of epithelial wound healing. LSCD can arise due to chemical burns, microbial infections, chronic inflammation, keratopathies, Stevens-Johnson syndrome, and graft-versus-host disease, often culminating in pain, scarring, and vision loss. Currently, transplantation remains the gold-standard therapeutic approach; however, it is limited by scarcity of donor tissue, requirement for regular follow ups and long-term immunosuppression, poor cell viability and variable graft survival. Consequently, alternative regenerative approaches, including stem cell-based therapies, de-cellularized matrices, tissue engineered constructs, exosomes and exogenous bioactive factors are being explored (Li et al., 2024). In this context, the proliferation of LESCs in response to DMSO+D3-induced HL-60 cells is encouraging, since enhancement of functional property of LESCs would be expected to produce better outcomes when used for healing of alkali-injured corneas.

Further, the observed regenerative effects of immature neutrophil-like population on scratch-injured HCEC cells and alkali-injured (*ex vivo* and *in vivo*) corneas can be attributed to the expression of key repair-associated molecular markers, such as arginase 1, annexin A5, galectin-1, S100A8 and S100A9. Among these markers, annexin A5 is known to promote corneal epithelial wound healing by promoting cell migration and tissue repair (Watanabe et al, 2006). S100 family proteins, particularly S100A4 and S100A9, play key roles in cellular differentiation, growth, maintenance of cellular architecture and regulatory processes occurring in the corneal limbus (Dua et al, 2005). Furthermore, arginase is reported to contribute to the immune privilege of the eye and is important for corneal graft survival (Fu et al, 2011). Galectins-3 and −7 compared to galectin-1 are known to be important for re-epithelialization of corneal wounds (Cao et al, 2002), while galectin-1 has been reported to be responsible for cutaneous wound healing (Lin et al, 2015). Its expression in these cells may therefore indirectly contribute to corneal repair.

The *in vivo* data demonstrated that DMSO+D3-induced immature neutrophil-like cells are useful for (i) effective restoration of the corneal epithelial layer, displaying increased expression of corneal epithelium specific marker KRT12; (ii) reduced corneal opacity, and attenuation of myofibroblast differentiation, as evidenced by decreased α-SMA expression and reduced fibronectin deposition; (iii) diminished neo-vessel formation; and (iv) modulation of the immune response, characterized by reduction in invasion of neutrophils and a transition towards a reparative M2 macrophage phenotype.

Collectively, these findings indicate that DMSO+D3-induced immature neutrophil-like cell population exhibit robust regenerative and immune-modulatory properties. Their consistent efficacy across *in vitro*, *ex vivo* and *in vivo* models highlight their potential as a novel and promising therapeutic strategy for corneal wound healing.

## Materials and methods

The study was approved by Institute Ethics Committee of PGIMER (INT/IEC/2022/SPL-1038).

### Cell culture and induction of differentiation to immature neutrophil-like cells

HL-60 cells (NCCS, Pune) were cultured in Roswell Park Memorial Institute 1640 (RPMI; Himedia, AT-150) medium, supplemented with 10% fetal bovine serum (FBS; Himedia, RM9955) and 40 µg/mL gentamycin (Himedia, TC026), and maintained at 37 °C in 5% CO_2_. For induction of differentiation into various immune cell types, HL-60 cells were seeded at a density of 1×10^5^ cells/mL in RPMI + 10% FBS and cultured with supplementation of (i) 1.3 % DMSO (Sigma-Aldrich, I10005), (ii) 100 nM 1α,25-dihydroxyvitamin D3 (D3; Abbott, Rocaltrol), (iii) 1.3% DMSO in combination with 100 nM D3. All cultures were maintained under these conditions for four days (unless otherwise specified) at 37 °C in 5% CO_2._

### Morphological analysis

Cells in each condition were examined for morphological changes, and phase contrast images were captured at 40X magnification using Leica microscope.

### Leishman staining

To assess the changes in nuclear morphology, Leishman staining was performed according to the manufacturer’s protocol (Himedia, S018S). Briefly, cells from the respective conditions were pooled, and a cell smear was prepared and rapidly air-dried, followed by the addition of Leishman stain for four minutes (methanol in the stain fixes the preparation). The stain was then diluted with double its volume of phosphate buffered saline (PBS) or distilled water (pH-7.0), gently mixed, and allowed to incubate for 10-12 minutes at room temperature. Slides were washed with PBS or distilled water and air dried. Images were captured at 40X magnification using a fluorescence microscope (EVOS FL Auto, Life technologies).

### cDNA synthesis and quantitative Real time PCR (qRT-PCR)

Total RNA from three-four biological replicates of cells cultured under different conditions was isolated using Trizol RNA isolation method (Sigma-Aldrich, TRI reagent, T9424) and from mouse corneal tissue was isolated using the ReliaPrep™ RNA Cell Miniprep System (Promega, Z6011; 50 preps). The concentration and purity of RNA was determined using NanoDrop spectrophotometer. cDNA was synthesized from 500-1000 ng RNA with Verso reverse transcriptase enzyme mix following manufacturer’s protocol (Thermo Scientific, AB-1453/A). Gene expression was quantified using SYBR green chemistry (Promega, GoTaq® qPCR, A6001) on a CFX96® Real-Time system (Bio-Rad). qRT-PCT was carried out in a final reaction volume of 10 µL, consisting of 5 µL GoTaq® qPCR Master Mix (2X), 1 µL cDNA (25 ng/µl), 1 µL gene-specific primers and 3 µL nuclease free water. Primer sequences used for each gene are listed in Table 2. PCR reaction for each target gene was performed in triplicates under the following thermal cycling conditions: 95°C for 10 minutes, followed by 40 cycles of 95°C for 45 s; 52, 57 or 60°C for 45 s; 72°C for 45 s. Microsoft Excel was used for data analysis after obtaining the cycle threshold (Ct) values from CFX Maestro software. For cells, gene expression was normalized with respect to the house-keeping gene, β-actin, and for mouse corneal tissue it was normalized to mouse 18S ribosomal RNA (18S rRNA) and expressed as fold change relative to controls using the 2−ΔΔCt method.

**Table 2:**
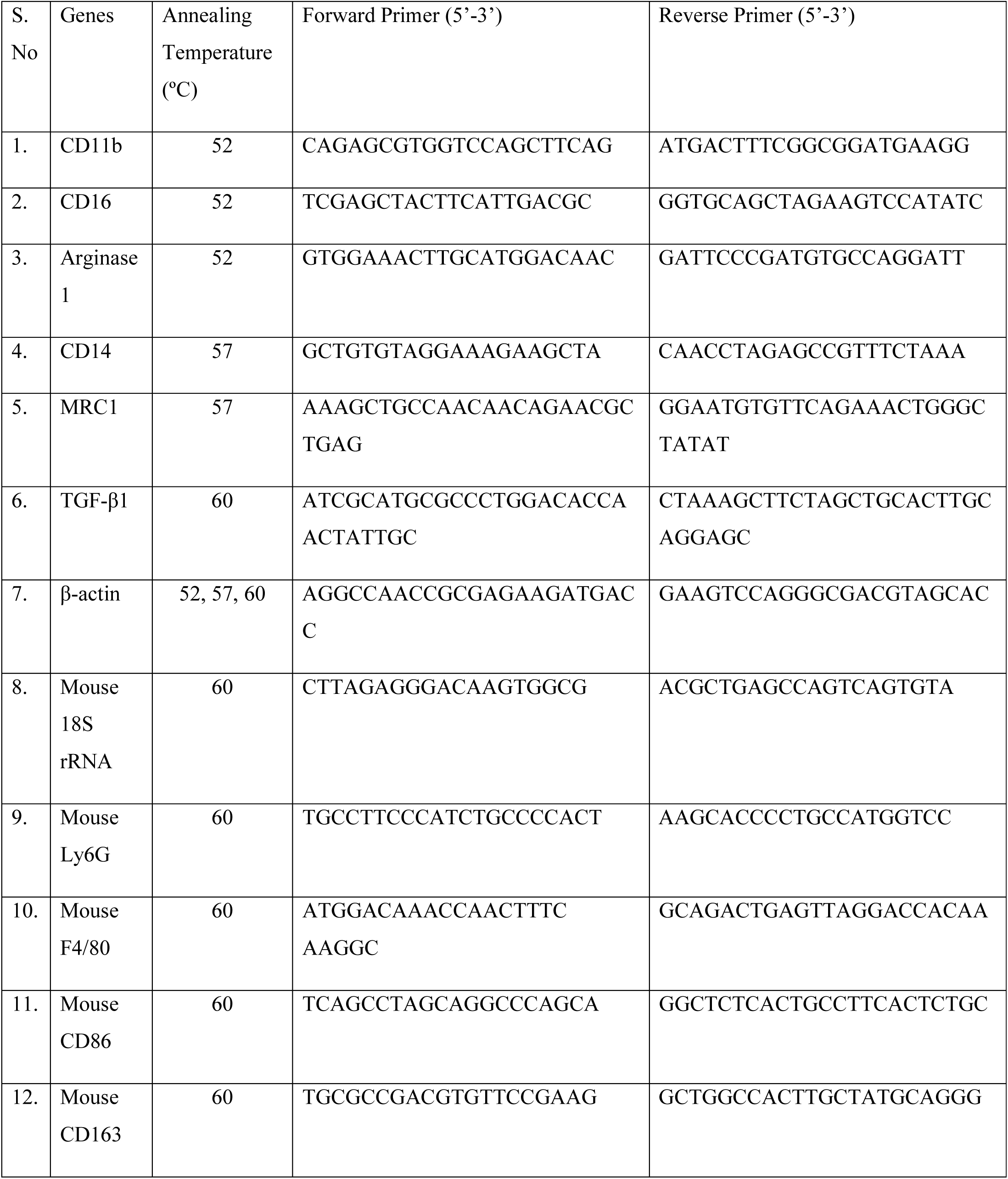
Primer sequences used for quantifying the expression of various markers by qRT-PCR.

### Flow cytometry

Expression of cell surface antigens (CD11b, CD16 and CD14) was analyzed by flow cytometry. The cells from each condition were pooled and washed with PBS, then stained with anti-human CD11b-PC7 (Beckman Coulter, A54822), anti-human CD16-PE (Beckman Coulter, A07766) and anti-human CD14-APC (Beckman Coulter, IM2580U) for 20 minutes at room temperature. After staining, cells were washed twice with PBS. Data acquisition was performed using a BD FACSCanto™II flow cytometer (BD Biosciences) and analyzed with BD FACSDiva 9.0 software.

### Measurement of Reactive oxygen species (ROS)

To assess ROS production, cells from each condition were divided into two tubes: (a) un-stimulated and (b) stimulated. Cells in both tubes were labelled with 0.5 mg/mL dihydrorhodamine 123 (DHR; Sigma-Aldrich, D1054) and incubated at 37°C for 15 minutes. Tube (b) was then stimulated with 10 ng/mL of phorbol-12-myristate-13-acetate (PMA; Sigma-Aldrich, P8139) for 15 minutes. After incubation, cells were washed with PBS, and data was acquired on a BD FACSCanto™II flow cytometer and analysed using BD FACSDiva 8.0.3 software. The neutrophil oxidative index (NOI) or stimulation index was calculated as the ratio of median fluorescence intensity in stimulated cells (DHR + PMA) to that in unstimulated cells (DHR only).

### Mass spectrometry proteomic analysis

Proteome analysis was performed using five biological replicates each of uninduced HL-60 cells and HL-60 cells-induced with DMSO+D3, following the protocol described by Thakur and Guptasarma (2022), with modifications in the lysis procedure. Cells were lysed in 90 µL of urea lysis buffer [7 M urea (MP, 194857), 2 M thiourea (Sigma-Aldrich, 62-56-6), 10 mM Tris-HCl (MP, 103133), pH-8.5] supplemented with 10 µL protease inhibitor (Sigma-Aldrich, P8340). The lysates were mixed by pipetting for 10 minutes on ice, sonicated for 2-3 seconds, incubated for 30 minutes, and centrifuged at 16,000 x g for 20 minutes at 4°C. Protein concentration in the cleared supernatants was determined using the Bicinchoninic acid (BCA) assay (GCC, Biotech). For trypsin digestion, 30 µg of protein per sample was processed as follows: reduced with 10 mM dithiothreitol (DTT; Sigma-Aldrich, D9779) for 1 h at room temperature, alkylated with 50 mM iodoacetamide (IAA; Sigma-Aldrich, I6125) for 1 h at room temperature and residual IAA quenched with 40 mM DTT for 1 h. Samples were diluted with 1 mM calcium chloride (CaCl_2_; Sigma-Aldrich, C7902, pH-7.6) to reduce the urea concentration and pH adjusted to 8-8.5 using 0.1 M Tris-Cl. Trypsin (Pierce™ Trypsin Protease, MS grade, 90057) was added at a concentration of 1 µg per sample and digestion was carried out for 12-16 h at 37°C. The reaction was stopped by lowering the pH to 2.0 using 10% trifluoroacetic acid (TFA; Thermo Fisher Scientific, A116-50). Sample clean-up was performed using C-18 resin columns (Thermo Scientific; Pierce™ C-18 spin columns, 89870), and vaccum dried, followed by reconstitution in solution containing 0.1 % formic acid (Fluka Analytical, 09676), 98 % LC-MS grade water (Pierce^TM^, 51104) and 2 % acetonitrile (ACN; Pierce^TM^, 51101). Mas spectrometric data was acquired in data-dependent mode on an Orbitrap Fusion Tribrid Mass Spectrometer (Thermo Fisher Scientific, Waltham, Mass., USA) connected to an EASY-nLC 1200 system, using the protocol described previously (Thakur and Guptasarma, 2022). The MS/MS data were deposited to the ProteomeXchange Consortium via the PRIDE (EMBL-EBI^©^, Cambridge, UK) partner repository with the dataset identifier PXD070913 and 10.6019/PXD070913. Proteome Discover 2.4 software (Thermo Fisher Scientific^©^) was used to analyze the raw files by searching against the Uniprot^©^ reference database; parameters used were: tolerance level, 10 ppm for MS and 0.05 Da for MS/MS; missed cleavages, 2; dynamic modifications, acetylation (protein N-term) and oxidation (M); fixed modifications, carbamidomethylation of cysteines.

For the calculation of fold change of individual proteins, the mean normalized abundance of each protein was calculated in the uninduced HL-60 group as well the DMSO+D3 group, followed by dividing the mean normalized abundance of a protein in HL-60-induced group vs the mean normalized abundance of that protein in the uninduced HL-60 group. Proteins were considered to be differentially expressed between the two groups, if the fold change was greater than or equal to 1.5 and false discovery rate-adjusted *P*-values was less than 0.05 (*P*<0.05). Reactome pathway analysis was performed using the Database for Annotation, Visualization, and Integrated Discovery (DAVID) software.

### Wound closure by scratch assay

The percentage of wound closure assessed by scratch assay was used as an *in vitro* model of corneal epithelial wound healing, with HCEC cells serving as the epithelial cell model. Cells were cultured in Dulbecco’s Modified Eagle Medium-Ham’s F12 (DMEM/F12) supplemented with 20 ng/mL epidermal growth factor (EGF; Sigma Aldrich, E9644), 4 µg/mL insulin (Biocon, Insugen-R), 10% fetal bovine serum and 40 µg/mL gentamicin and maintained at 37 °C and 5% CO_2_. HCEC cells were seeded in 24-well plates and cultured until confluence, followed by serum starvation for 6 h. A linear scratch injury was then created using a 10 μL plastic pipette tip, followed by a PBS wash to remove cell debris. HL-60 cells uninduced and induced with DMSO, DMSO + D3 and D3 (5× 10^5^ cells per insert) were seeded into the upper chamber of 0.4 μm culture inserts (SPL, 35324) and placed over the wounded HCEC in the bottom chamber of 24-well plate. Scratched HCEC cells without any direct contact with immune cells, but treated with serum-free medium (RPMI-SF), served as the control (Kacham et al, 2021). Migration of HCEC cells into the scratched area was monitored 24 h post-scratch injury, and images were captured. Image analysis and quantification of the percentage of wound closure (change in wound area over time) was performed using Wound_healing_size_tool_updated.ijm Image J plugin installed in Image J software.

### Colony formation assay

Human corneal limbal epithelial cells (HCLE) were used as an *in vitro* model for limbal epithelial stem cells (LESCs). These cells were cultured in keratinocyte serum-free medium (KSFM) supplemented with 30 µg/mL bovine pituitary extract (BPE), 0.2 ng/mL EGF, 0.3 mM calcium chloride (CaCl_2_) and 40 µg/mL gentamicin and maintained at 37 °C and 5% CO_2_. The potential of uninduced and induced HL-60 cells towards promoting colony formation of HCLE cells was evaluated following the method described by (Nam et al, 2017). HL-60 cells uninduced and induced with different reagents (5× 10^5^ cells/insert) were added to the upper chamber of 0.4 μm pore size culture inserts and co-cultured with HCLE cells (through an indirect contact) seeded at a density of 250 cells per well in 24 well plates (in bottom chamber) for seven days. At the end of the incubation period, colonies were fixed with 4% paraformaldehyde (PFA; Sigma Aldrich, P6148) for 15 minutes at 4°C and stained with 0.025% crystal violet (Himedia, TC510) prepared in 10% ethanol for 30 minutes, followed by washing twice with 1X PBS. Extraction of crystal violet dye was done using 0.2 M sodium dihydrogen phosphate (NaH2PO4; Himedia, GRM1255) diluted in ethanol (1:1) for 30 minutes. Quantitation was performed by measuring optical density at 570 nm. Images of the colonies were captured using a smartphone camera.

### *Ex vivo* model of corneal wound healing

An *ex vivo* goat alkali-injured corneal organ culture model was used to assess the ability of DMSO+D3-induced HL-60 cells for therapeutic effect in restoration of corneal homeostasis after chemical injury (Shukla et al., 2024). Briefly, goat eyes (n=3 per condition) were obtained from a local abattoir and transported in sterile PBS solution. All procedures were carried out in a laminar airflow hood. Excess surrounding tissue was removed from the goat eyeballs, which were then rinsed with betadine solution (Povidone-Iodine Solution IP, 10% w/v) and placed in high antibiotic PBS containing 5 µg/mL amphotericin B and 10X penicillin/streptomycin for 10 minutes. Corneal wound was created by application of 10 mm filter paper disk (cut using a 10-mm trephine) soaked in 1.0 M sodium hydroxide (NaOH) for 30 seconds on the corneal surface. Following injury, the wound was thoroughly washed with PBS to remove excess NaOH. The wounded eyeball was then held with tissue paper to provide support and pierced using a surgical blade, after which the corneal tissue was cut using surgical scissors to remove the remaining parts of the eye. A mounting medium consisting of DMEM/F12, 1 mg/ml bovine collagen (Advanced BioMatrix, 5005) and 1% agar (Himedia, GRM026P) was added to the opposite side of the corneal tissue. To maintain the corneas at an air-liquid interface, the corneal specimens were transferred to 6- or 12-well plates with the epithelial surface facing upward. 1 mL of organ culture medium consisting of DMEM/F12, 5% dextran (Himedia, T500), 1X penicillin/streptomycin and 2.5 µg/mL amphotericin B was added to each well. The corneas were then treated with uninduced HL-60 cells or induced with different reagents at a density of 5×10^5^ cells/eye daily and the cultures were maintained at 37°C and 5% CO2 for seven days. The extent of injury and wound healing was assessed by slit-lamp examination performed on days 0 and 7, while histological and immunofluorescence analysis were carried out at the termination of the experiment (day 7).

### *In vivo* model of corneal wound healing

An *in vivo* model of corneal wound repair was based on chemical burn induced in 6-8 weeks old, C57BL/6 female mice. The right eye of each mouse was alkali injured, while the contralateral left eye remained uninjured and served as a normal healthy control. All mice were anesthetized by intraperitoneal injection of 50 mg/kg ketamine (Ketapil®, Ketamine Injection I.P, 10 mL) and 5 mg/kg xylazine (XYLAXIN®, Xylazine Injection, 10 mL). In addition, local anesthesia was applied to the eye to be alkali injured, using 0.5% proparacaine hydrochloride ophthalmic solution (Paracain). Alkali injury was induced by placing a 2 mm filter paper disc (cut using a paper punch) soaked in 0.5 M NaOH, onto the corneal surface for 15 seconds, followed by thorough rinsing with 10 mL of 1X PBS to remove residual NaOH.

Mice were randomly divided into two groups: (i) vehicle and (ii) treatment. The injured right eye of vehicle group received fibrin glue alone, consisting of 1 μL fibrinogen (91 mg/mL; TISSEEL Lyo), followed by 0.5 μL thrombin (500 IU/mL, TISSEEL Lyo). In the treatment group, the injured right eye received 50 ×10^3^ DMSO+D3-induced HL-60 cells re-suspended in 1 μL fibrinogen, followed by 0.5 μL thrombin (Jhanji et al., 2022; Santra et al., 2024). To evaluate the efficacy of DMSO+D3-induced immature neutrophil-like cells, the vehicle group initially included n=8 mice, of which two died during the tenure of the experiment, while the treatment group included n=10 mice. Ophthalmic examinations were performed on days 0, 7 and 14, while histological, immunofluorescence and real-time PCR analysis were conducted at the end of the experiment (day 14).

### Ophthalmic examination

Clinical evaluations were carried out by an ophthalmologist (Advanced Eye Centre, PGIMER) using a slit-lamp microscope. The epithelial defects were evaluated using fluorescein strips (IO Floro) directly applied to the corneal surface. Fluorescein staining was examined under cobalt blue illumination, whereas bright-field images were captured under diffuse illumination of slit lamp microscope at 10X magnification for goat corneal tissues and 40X magnification for mouse corneal tissues. Imaging was performed at the initiation (day 0) and termination (day 7) of *ex vivo* experiment and days 0, 7 and 14 of the *in vivo* experiment.

Clinical scoring for in vivo experiment was performed based on the following criteria:

a. Corneal epithelial defect: Score 0, no defect; Score 1, defect less than 25%; Score 2, 25% to 50% epithelial defect; Score 3, 50% to 75% epithelial defect and; Score 4, more than 75% epithelial defect (Kim et al, 2022).
b. Corneal opacity: Score 0, completely clear; Score 1, slightly hazy with iris and pupil easily visible; Score 2, slightly opaque with iris and pupil still detectable; Score 3, opaque and pupil hardly detectable and; Score 4, completely opaque with no view of pupil (Anderson et al, 2014; Kim et al, 2022).
c. Neovascularization: Score 0, no neovessels; Score 1, neovessels at the corneal limbus; Score 2, neovessels spanning the corneal limbus and approaching the corneal center and; Score 3, neovessels spanning the corneal center (Anderson et al, 2014).

### Histological analysis

To analyse the histopathological changes, paraffin block preparation was carried out; corneal tissues fixed in 10% formalin, were dehydrated through graded ethanol series and were embedded in paraffin. Following paraffin block preparation, 3-4 µm thick sections were cut on the albumin coated slides. Hematoxylin and eosin (H&E) staining was performed.

### Immunofluorescence staining of tissue sections

Immunostaining of tissue sections was performed by following the protocol described by (Zaqout et al., 2020), with modifications in incubation durations. Tissue sections of 3-5 μm thickness were cut on 0.01% poly-L-lysine-coated slides for immunofluorescence analysis. Paraffin wax was melted by placing the slides at 80°C for 15 minutes, followed by further dewaxing using four changes of xylene (Fisher Scientific, 35417), with slides incubated for 5 minutes in each solution. This was followed by 5 minute incubation in xylene: ethanol (1:1) solution. The tissue sections were then rehydrated in a series of decreasing concentrations of ethanol solution (100%, 95%, 70% and 50%) for 5 minutes each, followed by washing in distilled water for 5 minutes. Antigen retrieval was performed using a heat-induced retrieval method with citrate buffer (pH-6.0). Citrate buffer was prepared by mixing Solution A and Solution B in a 9:1 ratio [Solution A: 0.1 M citric acid (Himedia, RM1023) in water; Solution B: 0.1 M tri-sodium citrate (Himedia, GRM3953) in water]. The citrate buffer was pre-heated in a microwave for 8 minutes while the slides were kept in water. Slides were then incubated for 3 minutes in the citrate buffer, followed by a wash with 1X PBS for 10 minutes. Permeabilization was carried out for 10 minutes using permeabilization solution containing 0.2% gelatin (Himedia, RM019) and 0.25% Triton-X-100 prepared in 1X PBS. Non-specific binding was blocked by incubating the sections with blocking solution (5% BSA in permeabilization solution) for 1 h at room temperature. Primary antibodies specific to the epithelial marker KRT12 (Santa Cruz Biotechnology INC, sc-515882) and fibrotic markers: α-SMA (Invitrogen, MA5-11547) and fibronectin (Invitrogen, 14-9869-82) were prepared in 1% BSA at a dilution of 1:100 and incubated overnight at 4 °C in a moist chamber. This was followed by one wash with 1X PBS for 5 minutes, and one with permeabilization solution for 5 minutes. Goat anti-mouse IgG Alexa Fluor^TM^ 488 (1:1000; Invitrogen, A11001) secondary antibody was then added and incubated for 1 h at 4 °C, followed by a wash with 1X PBS. Cell nuclei were stained with DAPI (1:10,000) for 15 minutes. Slides were subsequently washed with 1X PBS, wash solution [10 mM copper sulphate (CuSO_4_; Himedia, GRM677) and 50 mM ammonium chloride (NH_4_Cl; Himedia, RM3885) in water] and distilled water for 5-10 minutes each. Slides were air-dried and mounted using glycerol. Marker expression was analyzed by capturing images at 10X magnification for goat corneal tissues and 60X magnification for human and mouse corneal tissues using a confocal microscope (Olympus Fluoview FV3000).

## Statistical analysis

All data analyses were performed using Microsoft Excel (Microsoft Corporation, Redmond, WA, USA). Data are presented as mean ± standard deviation, and differences between groups were evaluated using Student’s t-test. A *P*-value of ≤ 0.05 was considered statistically significant (\**P* ≤ 0.05, \*\**P* < 0.01, \*\*\**P* < 0.001, ns: non-significant), where symbols indicate (*) increased and (#) decreased expression.

## Data availability

The mass spectrometry data has been deposited to the ProteomeXchange Consortium via the PRIDE repository with the dataset identifier PXD070913 and 10.6019/PXD070913.

## Author contributions SK

Data curation, Formal analysis, Validation, Investigation, Methodology, Writing-original draft, Writing-review and editing; **AS**: Assistance in in-vivo experimentation; **AG**: Visualization, Investigation; **BB**: Slit lamp examination; VS: Slit lamp examination; **UNS**: Visualization, Supervision, Investigation, Formal analysis; **PCG**: Visualization, Funding acquisition. **MLG**: Conceptualization; Resources; Supervision; Funding acquisition; Writing-original draft; Project administration; Writing-review and editing

## Acknowledgment

The authors thank Ms. Tanuja Kaushik and Ms. Meenakshi of the Central Sophisticated Instrument Cell (CSIC) for LCMS and confocal data acquisition respectively, and Mrs. Parveen and Ms. Aishwarya from Department of Hematology for assistance in flow cytometry experiments.

## Funding

This work was supported by an intramural grant from the PGIMER; grant no. IM/267/10-09-22-0455.

## Legends to Expanded view figures

**Figure EV1.** Flow cytometric gating strategy for the DHR assay and expression of surface markers in unstained uninduced HL-60 cells or upon induction with different reagents. Gating scheme for DHR assay is shown: (A) forward scatter (FSC) versus side scatter (SSC) and (B) scoring for the conversion of the non-fluorescent dihydrorhodamine (DHR) 123 to fluorescent rhodamine 123 (which emits green fluorescence), represented on the X-axis as median fluorescence intensity measured in the FITC channel, versus cell count on Y-axis. The gating scheme used for evaluation of expression of surface immune cell markers (CD11b, CD16 and CD14): (C) FSC versus SSC; (D) quadrant dot plot for scoring of APC (Y-axis) versus PE-Cy7 (X-axis) and (E) quadrant dot plot for scoring of PE-Cy7 (Y-axis) versus PE (X-axis) in unstained HL-60 cells, either uninduced or induced with DMSO + D3, DMSO alone or D3 alone, following induction for 4 days.

**Figure EV2.** Flow cytometric gating strategy for evaluation of surface marker expression at different time intervals in unstained uninduced HL-60 cells or upon induction with different reagents. The gating scheme used for evaluation of expression of surface immune cell markers (CD11b, CD16 and CD14): (A) FSC versus SSC and quadrant dot plot for scoring of (B) APC versus PE-Cy7 and (C) PE-Cy7 versus PE in unstained HL-60 cells, either uninduced or induced with DMSO and DMSO + D3 following induction for 4, 6 and 8 days.

**Figure EV3.** Expression of surface immune cell markers in HL-60 cells uninduced or upon induction with different reagents at different time intervals. Representative flow cytometric analysis showing the expression of (A) CD11b+CD14+ and (B) CD11b+CD16+ markers in uninduced HL-60 cells or upon induction with DMSO and DMSO + D3 for 4, 6 or 8 days respectively.

**Figure EV4.** Clinical evaluation of corneal healing following treatment of alkali-injured mice corneas (*in vivo*) with DMSO+D3-induced HL-60 cells or with vehicle alone. Quantitative comparison of (A) epithelial defect scores, (B) opacity scores and (C) neovascularization scores in alkali injured mice corneas treated with DMSO + D3-induced HL-60 cells compared with vehicle-treated (fibrin glue only) control corneas. Data are represented as mean ± standard deviation from one experiment with n=6 eyes in the vehicle group and n=10 eyes in the cells-treated group. Statistical significance was calculated using Student’s t-test; *P* ≤0.05 was considered significant [\**P* ≤ 0.05, \*\**P* < 0.01, \*\*\**P* < 0.001; symbols (*) and (#) indicate increased and decreased expression respectively].

